# Estimating the proportion of disease heritability mediated by gene expression levels

**DOI:** 10.1101/118018

**Authors:** Luke J. O’Connor, Alexander Gusev, Xuanyao Liu, Po-Ru Loh, Hilary K. Finucane, Alkes L. Price

**Affiliations:** Program in Bioinformatics and Integrative Genomics, Harvard Graduate School of Arts and Sciences, Cambridge, MA; Department of Epidemiology, Harvard T.H. Chan School of Public Health, Boston, MA; Dana Farber Cancer Institute, Boston, MA; Program in Medical and Population Genetics, Broad Institute, Cambridge, MA; Department of Mathematics, Massachusetts Institute of Technology, Cambridge, MA; Department of Biostatistics, Harvard T.H. Chan School of Public Health, Boston, MA

**Author notes:** Correspondence should be addressed to L.J.O or A.L.P.

## Abstract

Disease risk variants identified by GWAS are predominantly noncoding, suggesting that gene regulation plays an important role. eQTL studies in unaffected individuals are often used to link disease-associated variants with the genes they regulate, relying on the hypothesis that noncoding regulatory effects are mediated by steady-state expression levels. To test this hypothesis, we developed a method to estimate the proportion of disease heritability mediated by the cis-genetic component of assayed gene expression levels. The method, gene expression co-score regression (GECS regression), relies on the idea that, for a gene whose expression level affects a phenotype, SNPs with similar effects on the expression of that gene will have similar phenotypic effects. In order to distinguish directional effects mediated by gene expression from non-directional pleiotropic or tagging effects, GECS regression operates on pairs of cis SNPs in linkage equilibrium, regressing pairwise products of disease effect sizes on products of cis-eQTL effect sizes. We verified that GECS regression produces robust estimates of mediated effects in simulations. We applied the method to eQTL data in 44 tissues from the GTEx consortium (average *N*_eQTL_ = 158 samples) in conjunction with GWAS summary statistics for 30 diseases and complex traits (average *N_GWAS_* = *88K*) with low pairwise genetic correlation, estimating the proportion of SNP-heritability mediated by the cis-genetic component of assayed gene expression in the union of the 44 tissues. The mean estimate was 0.21 (s.e. = 0.01) across 30 traits, with a significantly positive estimate *(p <* 0.001) for every trait. Thus, assayed gene expression in bulk tissues mediates a statistically significant but modest proportion of disease heritability, motivating the development of additional assays to capture regulatory effects and the use of our method to estimate how much disease heritability they mediate.

## Introduction

In the past few years, large scale expression quantitative trait locus (eQTL) studies have identified genetic variants associated with the expression levels of thousands of genes [1–6], motivated by the observation that disease risk variants identified by genome-wide association studies (GWAS) are predominantly noncoding [7]. In addition, GWAS risk variants and SNP-heritability (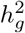; [8,9]) are enriched near epigenetic marks associated with regulatory function [10–15] and at eQTLs [16–20,22]. Cis-genetic correlations have been identified between complex traits and gene expression levels of specific genes [23–26], suggesting that eQTL studies can elucidate regulatory disease mechanisms.

However, the interpretation of observed enrichments and cis-genetic correlations between eQTLs and disease risk variants remains in doubt. Enrichment of GWAS signals near eQTLs could arise either from mediation of genetic effects by gene expression levels or from coincidental colocalization of eQTLs and GWAS variants, either due to pleiotropic effects of the same causal SNP or due to tagging effects arising from linkage disequilibrium (LD) between distinct causal SNPs. Indeed, most instances of colocalization between significant autoimmune disease loci and eQTLs are driven by tagging effects, suggesting that coincidental colocalization may be more prevalent than mediation [27]. Cis-genetic correlations can be primarily driven by a single large cis eQTL, so these correlations might also reflect coincidental colocalization [24]. The interpretation of enrichment and cis-genetic correlations depends on the extent that true mediation exists across the genome: if it is prevalent, then a mechanistic basis for these observations is plausible.

Here we develop a method, gene expression co-score regression (GECS regression), to estimate the proportion of disease heritability that is mediated by the cis-genetic component of assayed gene expression. In order to distinguish directional effects mediated by gene expression from non-directional pleiotropic or tagging effects, GECS regression operates on unlinked pairs of SNPs with shared cis genes. It is based on the intuition that if phenotypic effects are mediated by gene expression, then SNPs with similar effects on expression will have similar effects on the phenotype. Because GECS regression operates on GWAS summary statistics, it is broadly applicable to large publicly available data sets [28]. We applied our method to eQTL data in 44 tissues from the GTEx consortium [3] (see URLs) and GWAS summary statistics for 30 genetically distinct complex traits.

## Results

### Overview of methods

We aim to estimate the proportion of heritability mediated by the cis-genetic component of assayed gene expression levels. In particular, we model *underlying* (random-effect) cis-genetic correlations, which reflect true biological effects, instead of fixed-effect genetic correlations; the latter may be inflated by coincidental colocalization between causal effects (see Online Methods). Our definition of mediation does not precisely correspond to direct causality, as assayed expression may act as an imperfect proxy for truly causal expression levels in some other cell type or cellular context; however, it is unlikely to reflect *reverse* causality (see Online Methods and Supplementary Note). In order to estimate the proportion of heritability mediated by expression, we use an approach that is based on pairs of unlinked SNPs. Specifically, we regress pairwise products of phenotypic effects on products of cis-eQTL effect sizes. Other approaches are unable to recover this parameter, instead allowing contamination from non-mediated effects (see Supplementary Note). On the other hand, our method is not contaminated by non-mediated effects because products of non-mediated phenotypic effects are uncorrelated with products of eQTL effects for unlinked SNPs.

Under a random effects model, the expected value of the product between the marginal trait effect sizes *a_i_* and *a_j_* for a pair of unlinked SNPs (*i, j*) is:

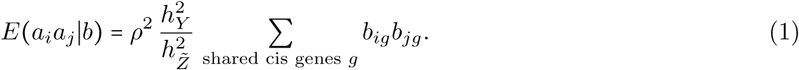

Here *ρ*^2^ is the proportion of heritability mediated by expression, 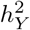 is the trait SNP-heritability, 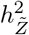 is the mean cis-heritability of expression levels, and *b_ig_* is the marginal effect of SNP *i* on gene *g* (see Online Methods and Supplementary Note). We refer to the sum over genes of products of cis-eQTL effect sizes as the “GE co-score” of SNPs *i* and *j*. Equation (1) does not hold for SNPs in LD; in the presence of positive LD, both *a_i_a_j_* and Σ*b_ig_b_jg_* will be positively inflated due to tagging of non-mediated effects. Equation (1) also relies on an independence assumption between the trait effect size and cis-genetic architecture of each gene (see Online Methods).

Equation (1) suggests an inference method for *ρ*^2^: we regress products of estimated trait effect sizes on estimated GE co-scores for pairs of unlinked SNPs. We refer to this method as GE co-score regression (GECS regression). We define unlinked SNPs using an LD threshold, retaining 75% of SNP pairs that have shared cis genes (with LD computed in the gene expression cohort), and we additionally condition the regression on LD for retained SNP pairs. We use eQTL data from multiple tissues; genes in each tissue are pooled, resulting in an estimate of the heritability mediated by the union of tissues. Because cis-genetic correlations are stronger than environmental correlations across tissues, this approach increases the effective cis-heritability, resulting in increased power (see Online Methods and Supplementary Note). We employ a weighted regression to further increase power. To avoid attenuation bias, we normalize the regression slope using a cross-validation estimate of the variance in true (rather than estimated) GE co-scores. The regression slope is multiplied by the ratio 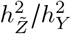 of expression heritability to trait heritability, estimated using LD score regression [34] (see URLs), to produce an estimate of *ρ*^2^. We assess standard errors using a jackknife on 50 contiguous blocks of SNPs, analagous to previous work [34]. To assess statistical significance, we jackknife only the regression numerator, and apply a *Z*-test. Further details of the method are provided in Online Methods. We have released open source software implementing the method (see URLs).

### Simulations

To assess the robustness of our methods, we performed simulations using a subset of real genotypes from the UK Biobank [44] (see URLs) (*N*_GWAS_ = 10,000; *M =* 5,000 SNPs) and simulated complex trait and gene expression phenotypes (N_eQTL_ = 1000 samples, disjoint from the GWAS samples; 600 genes; 50 tissues). We sampled effects of SNPs on expression, effects of expression on trait, and direct (non-mediated) effects of SNPs on trait from point-normal distributions. We included strong genetic and environmental correlations across tissues. To model coincidental colocalization between eQTLs and non-mediated GWAS variants, SNPs with a large effect on gene expression were more likely to have a nonzero direct effect on the trait (see Online Methods).

We ran GECS regression on the simulated data. When LD pruning was appropriately stringent, our estimates of *ρ*^2^ were approximately unbiased (Figure 1-a). Without LD pruning, more significant inflation due to LD was observed (Figure 1-b). At larger values of *ρ*^2^, modest downwards bias was observed at stringent LD thresholds, due to bias in gene expression heritability estimates obtained using LD score regression. This bias occurs because regression SNPs in the cis window are in LD with non-causal SNPs outside the cis window, which contribute to the LD score of the regression SNP but not to its χ^2^ statistic; the amount of bias is small (see Supplementary Note).

A naive fixed-effect estimator of *ρ*^2^ would be the sum of the squares, over genes, of the estimated fixed-effect cis-genetic correlation between that gene’s normalized expression level and the trait. To estimate the fixed-effect cis-genetic correlation, we took the dot product between the effect size estimates for each trait and divided by the geometric mean of the true simulated heritability for expression and trait. We evaluated the fixed-effect estimator in simplified simulations with no LD and no correlations among genes (see Online Methods). We hypothesized that, even in the absence of mediation, fixed-effect genetic correlations would arise by chance due to coincidental colocalization between eQTLs and non-mediated effects. We estimated the fixed-effect genetic covariance between each gene and the trait using the covariance between the effect size estimates, summed the squared estimates, and divided by the trait heritability (see Online Methods). Estimates of *ρ*^2^ using this method were close to 1, even when N_eQTL_ and *N*_GWAS_ were large (so sampling noise did not contribute to the bias). GECS regression estimates, on the other hand, were unbiased (Figure 1-c). We note that in the special case of no LD, our nave fixed-effect estimator is equivalent to cross-trait LD score regression with constrained intercept [35] and equivalent to *ρ*-HESS [29,30], implying that those methods are similarly impacted by coincidental colocalization. We are not currently aware of a method that provides an unbiased estimate of the squared random-effect genetic correlation under a sparse genetic architecture.

A more sophisticated approach is to partition trait heritability using eQTLs as an annotation [20,22], comparing the heritability explained by cis-eQTL SNPs with the heritability explained by other SNPs. We called the most significant 25% of SNPs as cis-eQTLs (similar to the proportion of SNPs that were called as eQTLs in [20,22]) and tested for enrichment of trait heritability explained by those SNPs using stratified LD score regression [14] (see Online Methods). Because our simulations include colocalization between effects on expression and direct (non-mediated) phenotypic effects, stratified LD score regression detects enrichment even in the absence of mediation (Figure 1-d)); this does not imply that stratified LD score regression or other heritability partitioning methods are flawed, but does imply that they cannot specifically distinguish mediation from colocalization effects. On the other hand, GECS regression had a well-calibrated falsepositive rate at *ρ*^2^ = 0, with increasing power to detect mediation as *ρ*^2^ increased. We caution that power may depend heavily on the cis-genetic architecture of cis-expression and the genetic architecture of the trait.

We assessed the calibration of our jackknife standard errors and *p* values. We normalized the empirical errors (the signed differences between the true and estimated values) by dividing them by the jackknife standard errors; the standard deviation of the normalized errors was slightly smaller than 1 on average, indicating that estimated standard errors are modestly conservative (Figure S1; Online Methods). When we calculated p values using a *Z* test, we found that they were conservative under the null (2% positive rate at *α =* 0.05, Figure 1-d).

**Figure 1:**
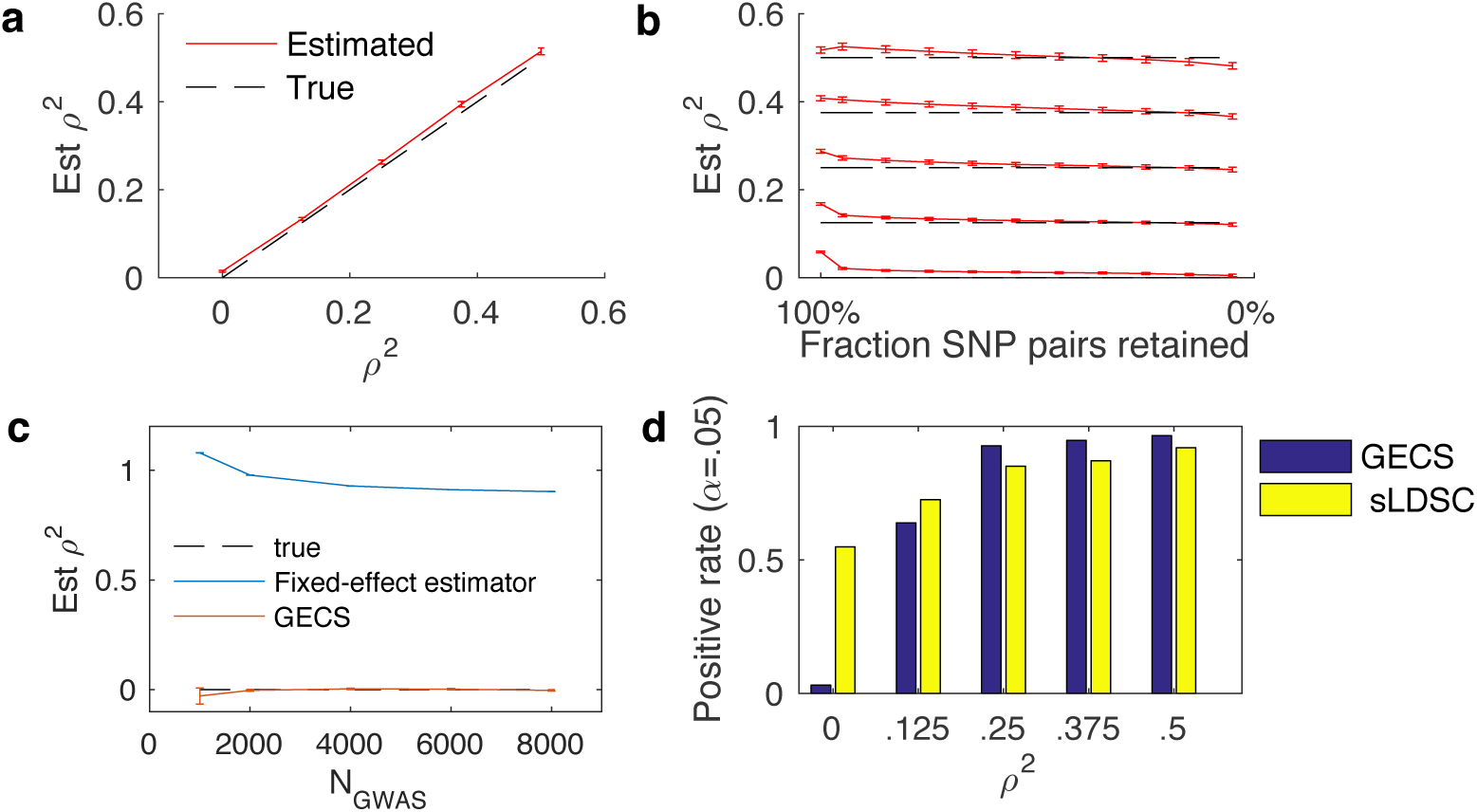
Performance of GE score regression in simulations using real genotypes and simulated gene expression and phenotypes. Simulations included strong colocalization between eQTLs and non-mediated disease SNPs. (a) Estimates were approximately unbiased when 75% of SNP pairs were retained. (b) Estimates were inflated when no LD pruning was applied, but LD pruning reduced the bias. Dashed horizontal lines indicate true values of *ρ*^2^. (c) Estimates of *ρ^2^* using GECS regression and a naive fixed-effect estimator at different eQTL and GWAS sample sizes, in a simplified simulation setup including colocalization but no mediation. (d) Comparison of power to detect an effect between GECS regression and stratified LD score regression (sLDSC), using eQTLs as an annotation. Error bars indicate standard error based on 288 simulations (a,b,d) or 40,000 simulations (c). Numerical results are reported in Table S1.

We investigated the impact on our estimates of varying the cis window size. If the cis window is too small, there are two possible sources of bias. First, downwards bias is introduced because cis-heritable signal is missed. Second, genetic correlations between nearby genes, in conjunction with LD between SNPs that are in cis for each gene, lead to upwards bias due to over-counting; this occurs specifically in the presence of genetic correlations between adjacent genes when the cis window is too small (see Supplementary Note). We varied the GECS regression cis window size while fixing the true cis window size. When the GECS regression cis window size was smaller than the true cis window size, cis-heritable signal was missed, and the estimator was downwardly biased (Figure S2-a). However, estimates stabilized when the GECS regression cis window was as large as the true cis window size (±1Mb). In the presence of correlations between nearby genes (as opposed to only across tissues), estimates of heritability explained per unit of cis-heritability were upwardly biased when GECS regression cis window was smaller than ±1Mb (Figure S2-b).

### Application to 30 complex traits

We applied GECS regression to summary statistics for 30 genetically distinct diseases and complex traits (Table S2; Online Methods). These included publicly available summary statistics from large consortia and summary statistics computed from UK Biobank data [44]. UK Biobank summary statistics were computed by applying BOLT-LMM [31] to up to 145,416 unrelated European-ancestry samples. We used gene expression data in 44 tissues from the GTEx consortium [3] (Table S3), computing cis eQTL summary statistics using FastQTL [45]. Expression summary statistics were computed from unrelated individuals consistent with GTEx consortium methods, and we additionally restricted to European samples (85% of individuals).

Estimates of the proportion of trait heritability mediated by the cis-genetic component of assayed gene expression are displayed in Figure 2 and Table 1. The mean estimate was 0.21 (s.e. = 0.01). Estimates were significantly positive (*p* < 0.001) for every trait. Except for multiple sclerosis, which had a large point estimates and a large standard error, estimates of *ρ*^2^ ranged between 0.07 and 0.41. The variation in *ρ*^2^ estimates was due at least in part to differences between phenotypes, rather than sampling error. The variance in *ρ*^2^ estimates was 0.013, compared with a mean squared jackknife standard error of 0.009 (which is likely to be a slight overestimate; Figure S1). A conservative estimate of the variance in true values of *ρ*^2^ across phenotypes, not accounting for positive correlations between noise terms, is the difference between these values, giving a standard deviation of 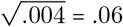 (31% of the mean). Noise terms are more likely to be positively correlated than negatively correlated, as a significant portion of the error comes from estimation of the regression denominator (the variance in GE co-scores), and this portion is highly positively correlated across phenotypes. This value 0.06 is statistically significant, again under the conservative assumption that sampling errors are uncorrelated (*χ*^2^ test *p* < 10^−10^; see Online Methods). On average, phenotypes with larger estimates also had larger standard errors. This phenomenon arises from the fact that variance in the regression denominator represents a significant portion of the sampling error, while the regression numerator (covariance) contains a large percentage of the variation across phenotypes; therefore, much of the observed error represents a percentage of the point estimate. This phenomenon also explains why estimates can be highly significantly positive (using the numerator as the test statistic) despite relatively large standard errors (Figure 2 and Table 1). Standard errors were also larger for phenotypes with less significant heritability estimates, with a Spearman correlation of 0.40 (*p =* 0.02) between the significance of 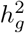 and the significance of *ρ*^2^.

**Figure 2:**
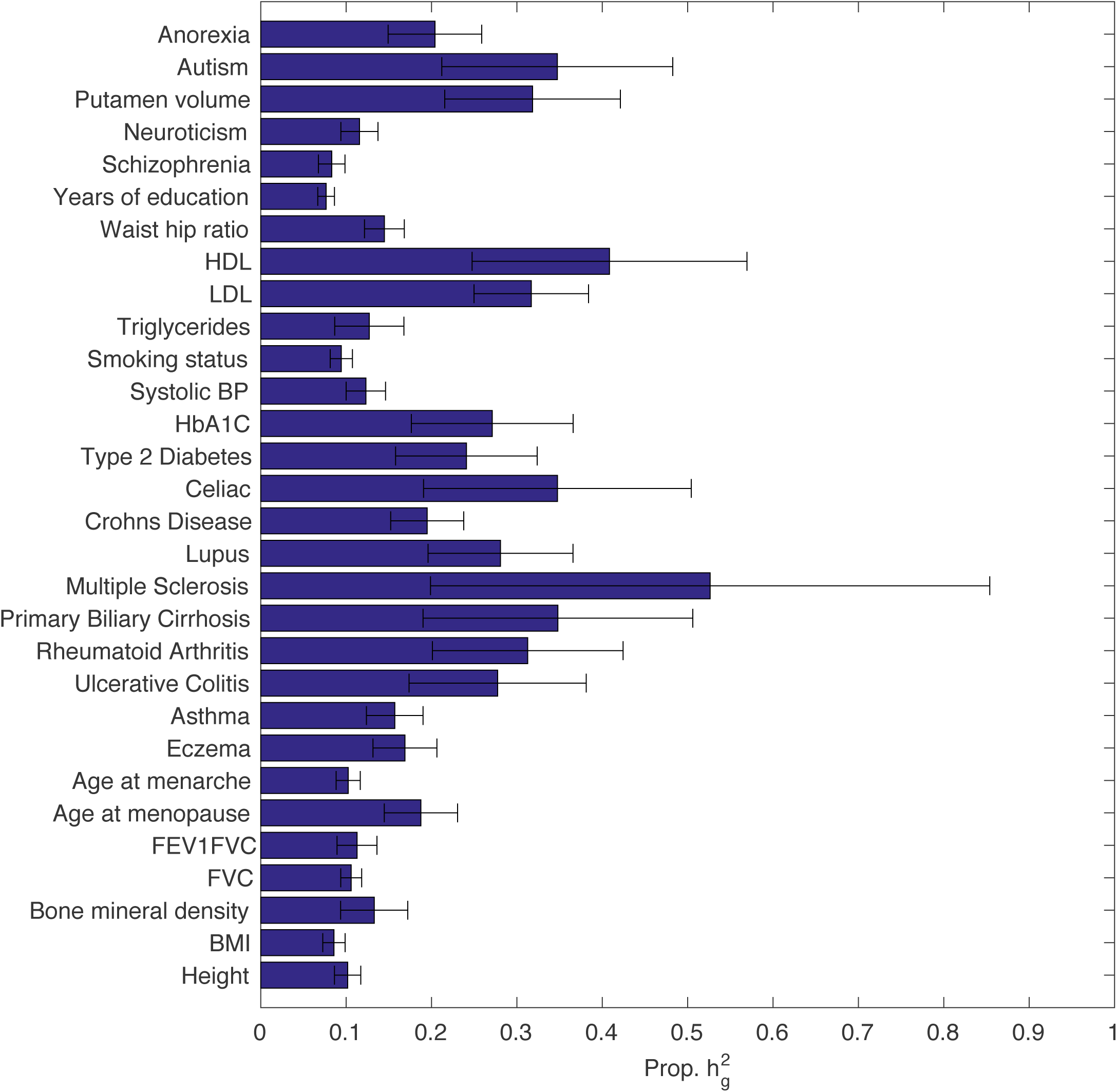
Estimates of the proportion of trait heritability mediated by the cis-genetic component of assayed gene expression for 30 diseases and complex traits. Related traits are grouped, and order is alphabetical within groups. Error bars represent jackknife standard errors. Numerical results are reported in Table 1.

**Table 1:** Estimates of heritability mediated by expression for 30 genetically distinct phenotypes. We report point estimates of *ρ*^2^ and 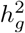 for each trait, together with standard errors and significance of *ρ* ^2^ estimates.

| Phenotype | $\rho^2$ | std err | Z score | $h_g^2$ | std err |
| --- | --- | --- | --- | --- | --- |
| Anorexia | 0.2 | 0.05 | 8.3 | 0.2 | 0.03 |
| Autism spectrum | 0.35 | 0.14 | 11.5 | 0.39 | 0.08 |
| Putamen volume | 0.32 | 0.1 | 8.8 | 0.37 | 0.08 |
| Neuroticism | 0.12 | 0.02 | 11.6 | 0.1 | 0.01 |
| Schizophrenia | 0.08 | 0.02 | 11.2 | 0.5 | 0.03 |
| Years of Education | 0.08 | 0.01 | 8.9 | 0.13 | 0.01 |
| Waist-hip ratio (BMI adjusted) | 0.14 | 0.02 | 7.5 | 0.14 | 0.02 |
| HDL | 0.41 | 0.16 | 3.7 | 0.06 | 0.03 |
| LDL | 0.32 | 0.07 | 5.4 | 0.09 | 0.03 |
| Triglycerides | 0.13 | 0.04 | 9.6 | 0.15 | 0.04 |
| Smoking status | 0.09 | 0.01 | 11.9 | 0.14 | 0.01 |
| Blood pressue (Systolic) | 0.12 | 0.02 | 11.8 | 0.2 | 0.01 |
| HbA1C | 0.27 | 0.09 | 5.4 | 0.1 | 0.02 |
| Type 2 Diabetes | 0.24 | 0.08 | 7.3 | 0.09 | 0.02 |
| Celiac Disease | 0.35 | 0.16 | 6.9 | 0.3 | 0.1 |
| Crohns Disease | 0.19 | 0.04 | 8.8 | 0.51 | 0.1 |
| Lupus | 0.28 | 0.08 | 6.3 | 0.43 | 0.1 |
| Multiple Sclerosis | 0.53 | 0.33 | 5.8 | 0.1 | 0.05 |
| Primary Biliary Cirrhosis | 0.35 | 0.16 | 9 | 0.46 | 0.11 |
| Rheumatoid Arthritis | 0.31 | 0.11 | 8.6 | 0.15 | 0.04 |
| Ulcerative Colitis | 0.28 | 0.1 | 9.3 | 0.24 | 0.05 |
| Asthma | 0.16 | 0.03 | 8.7 | 0.07 | 0.01 |
| Eczema | 0.17 | 0.04 | 12.3 | 0.08 | 0.01 |
| Age at menarche | 0.1 | 0.01 | 6.8 | 0.3 | 0.02 |
| Age at menopause | 0.19 | 0.04 | 10.1 | 0.19 | 0.03 |
| Lung FEV1/FVC ratio | 0.11 | 0.02 | 11.9 | 0.25 | 0.02 |
| Lung forced expiratory volume (FVC) | 0.11 | 0.01 | 10.4 | 0.25 | 0.02 |
| Bone mineral density (heel) | 0.13 | 0.04 | 5.8 | 0.29 | 0.03 |
| BMI | 0.09 | 0.01 | 8.8 | 0.3 | 0.02 |
| Height | 0.1 | 0.02 | 9.5 | 0.53 | 0.04 |

To assess the robustness of our estimates, we performed five secondary analyses. First, we varied our LD pruning threshold, finding that estimates were stable when we pruned stringently (Figure S3). We also investigated different LD pruning schemes. GECS regression prunes SNP pairs using a composite ranking of estimated LD and estimated *shared* LD (see Online Methods); however, concordant estimates were obtained when pruning using LD only or shared LD only, or when the regression was not conditioned on estimated LD (Table 2). Second, we investigated the impact of our regression weighting scheme, which down-weights SNP pairs whose SNPs have high LD to other regression SNPs; estimates using the weighted regression were concordant with estimates from the unweighted regression (Table 2). Estimates were also concordant if we removed the MAF lower bound (2*f* (1 − *f*) > .05), or if we used a stringent lower bound (2*f* (1 − *f*) > .25; Table 2). Third, to investigate the possibility that we were missing cis-genetic signal by restricting to a cis window of ±1Mb for each gene, we varied the cis window size. Estimates did not increase when we increased the window size to ±2Mb, indicating that little cis-genetic signal is missed (Figure S4). When we reduced the cis window size, we observed an increase in heritability explained and in heritability explained per unit of cis heritability, probably due to LD between cis windows for genetically correlated genes. This phenomenon, which is only expected to occur when significant heritability localizes to the flanking regions of genes, was also observed to a lesser extent in simulations (Figure S2; see Supplementary Note). Fourth, to determine whether the results were dependent on the sample size in the eQTL data set, we split the eQTL tissues into large N and small N subsamples (with 14 and 30 tissues, respectively). Estimates from these subsamples were concordant (Table 2). We also investigated the impact of randomly reducing the number of tissues (Figure S5). Estimates were relatively stable when ≥ 10 tissues were used; standard errors were particularly large when only 1 tissue was used. Fifth, we compared *ρ*^2^ estimates for pairs of summary statistic data sets that were pruned from the primary analysis due to high genetic correlation between traits (Table S4). Most estimates were concordant. Nominally significant differences were observed for 4 out of 15 pairs of traits; only one of these differences (*p* = . 002 for the two Years of Education datasets, which are overlapping) was significant after correcting for 15 pairs tested.

**Table 2:** Estimated heritability mediated by expression for different modifications to GECS regression, on average across non-redundant phenotypes. Standard errors were estimated by jackknifing. Different MAF ranges were used for estimation of GECS regression slope, but the same SNP set (heterozygosity ≥ .05) was retained for heritability estimation.

| | Mean estimated $\rho^2$ | Standard error |
| --- | --- | --- |
| Default parameters | .21 | .01 |
| Pruning on LD only | .21 | .01 |
| Pruning on shared LD only | .21 | .02 |
| No LD conditioning | .23 | .03 |
| No regression weighting | .22 | .02 |
| No MAF lower bound | .21 | .01 |
| Stringent MAF lower bound | .21 | .01 |
| Large $N_{\text{eQTL}}$ tissues | .25 | .01 |
| Small $N_{\text{eQTL}}$ tissues | .22 | .01 |

Finally, to investigate whether mediation of disease heritability by gene expression levels is tissue-specific, we estimated the proportion of heritability mediated by expression in 13 brain-related tissues, including 10 brain regions and tibial nerve, thyroid, and pituitary gland (Table S3). We assessed the heritability mediated by expression in both brain-related and non-brain-related tissues across nine brain-related traits: anorexia, autism, putamen volume, neuroticism, schizophrenia, years of education, ever smoked, age at menarche and BMI. Averaging across the nine traits, no difference was detected between heritability mediated by brain-related and non-brain-related tissues (*p* = 0.76; Table 3). Instead, both collections of tissues captured the full signal observed in their union (*p* = 0.69 and *p* = 0.49 respectively for the difference between heritability mediated by brain-related tissues and non-brain-related tissues compared with all tissues). These results are consistent with shared cis-eQTL effects and strong cis-genetic correlations across tissues [3, 36, 37].

**Table 3:** Estimated heritability mediated by expression for nine brain-related traits: anorexia, autism, putamen volume, neuroticism, schizophrenia, years of education, ever smoked, age at menarche and BMI. There was no significant difference between any of the three sets of tissues.

| | Mean estimated $\rho^2$ | Standard error |
| --- | --- | --- |
| Brain-related tissues and cognitive-related traits | .17 | .02 |
| Non-brain-related tissues and cognitive-related traits | .18 | .02 |
| All tissues and cognitive-related traits | .17 | .01 |

## Discussion

We have introduced GE co-score regression (GECS regression), a method to estimate the proportion of disease heritability mediated by gene expression levels; we have shown that the method can distinguish between mediation and coincidental colocalization effects, unlike other methods. We applied GECS regression to eQTL data in 44 tissues and GWAS summary statistics for 30 genetically distinct diseases and complex traits. We determined that the cis-genetic component of gene expression in the union of these tissues mediates a modest proportion of SNP-heritability, with a mean estimate of 0.21 (s.e. = 0.01). We observed no evidence of substantial tissue-specific cis effects, consistent with previous work [3,36,37]. On the one hand, these results provide validation that eQTL studies do capture regulatory disease mechanisms; on the other hand, they motivate the development of new assays to systematically identify the transcriptional targets of regulatory GWAS variants.

Which other regulatory effects might influence disease? First, some disease heritability may be mediated through mechanisms that could be captured using gene expression data but that were not considered in this study, such as alternative splicing or trans regulatory effects. We anticipate that upcoming GTEx data will include alternative splicing phenotypes, and GECS regression could be applied to this data; on the other hand, larger sample sizes will be needed before GECS regression will produce well-powered estimates using trans eQTL data. Second, regulation might occur in specific cell types or in response to extracellular stimuli, such as glucose availability or immune activation [21]. If this is the case, then eQTL studies in specific cell types under experimental conditions mimicking physiologically relevant contexts would increase heritability explained. Even if cell type-and context-specific data is collected at modest sample sizes initially, GECS regression could be applied to this data together with GTEx data in order to quantify the additional contribution of cell type-and context-specific effects. Third, regulation might be dynamic and transient, occurring sporadically in individual cells in response to internal states that are not synchronized across cells in a tissue. This type of effect might be detected using single-cell eQTL studies in conjunction with sophisticated statistical methods that identify the subset of cells that are in the disease-relevant state, or using hQTL and meQTL data [39–43] if epigenetic changes accompanying transient regulatory events are more stable. Similar to context-specific effects, the proportion of additional heritability mediated by this data could be estimated using GECS regression.

Our approach has several limitations. First, our method requires the assumption that genes with multiple independent eQTLs do not have systematically larger or smaller effect sizes than genes with only one eQTL. Second, our method requires that intermediate phenotypes often have multiple independent QTLs; for molecular phenotypes that rarely have more than one QTL, our method will produce noisy estimates, although the estimates will remain unbiased. Third, our method does not identify which genes are responsible for the signal that is detected; existing methods [23–26] have been used to identify these genes based on fixed-effect genetic correlations, but our results motivate the development of methods testing for the presence of random-effect genetic correlations with individual genes to avoid signals due to coincidental colocalization. Fourth, our method requires large effective eQTL sample sizes, which we have achieved by combining eQTL data across tissues (Figure S5). Fifth, as noted above, power limitations prevent us from applying GECS regression to the trans-genetic component of expression levels, although our estimates will include trans effects that are mediated by cis effects on different genes [38]. Sixth, it would be interesting to quantify the extent to which trans regulation is mediated by cis effects, by using trans expression data instead of disease data, but our method currently requires disjoint samples in the eQTL and GWAS cohorts. Seventh, our method is limited to additive effects, and an entirely different approach would be needed in order to quantify non-additive effects.

In conclusion, we have determined that the cis-genetic component of gene expression mediates a modest proportion of disease and complex trait heritability. These results motivate the continued development of new molecular QTL studies, and the use of GECS regression to quantify their improvement over existing assays in capturing regulatory effects that contribute to human disease.

## Acknowledgements

We are grateful to Soumya Raychaudhuri for helpful discussions and to Steven Gazal for assistance with GWAS summary statistics. This research was funded by NIH grants T32 GM074897, T32 HG002295, R01 MH107649, R01 MH101244 and U01 HG009379.

## URLs

GE co-score regression software will publicly released prior to publication at http://github.com/lukejoconn**or/** <u>gecs</u>. Description of GTEx data and publicly available data, http://gtexportal.org [3]; LD score re-gression software, http://github.com/bulik/ldsc [14,34,35]; description of UK Biobank data, http://www.ukbiobank.ac.uk/ [44].

## Online Methods

### Definition of heritability mediated by expression

We model a quantitative trait *Y*:

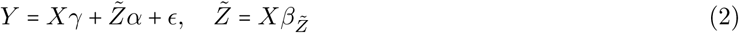
where *X* is the random genotype vector, *γ* is the random vector of direct SNP effects on the trait, *Z̃* is the cis-genetic component of gene expression, *α* is the random vector of gene effects on the trait, e is the environmental component of *Y*, and *β_Z̃_* is the random M × G matrix of cis-eQTL effects. Effects of the transgenetic component of expression are incorporated into *X_γ_*, and effects of the environmental component of expression are incorporated into *ϵ*. We do not assume that SNP and gene effect sizes are drawn from a specific family of distributions. The proportion of heritability mediated by expression is defined as

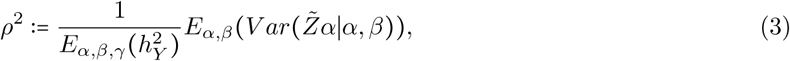

where 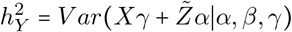 is the SNP-heritability of Y.

We use the term *mediation* because random-effect cis-genetic correlations require mechanistic rather than stochastic explanations. In particular, the random-effects parameter *ρ*^2^ differs from the expected maximum prediction accuracy of the genetic component of Y from *Z̃ fixing γ*:

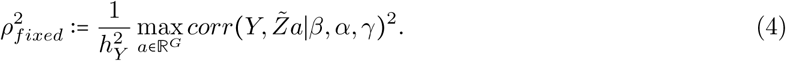

The arg max above is not equal to α, since coincidental genetic correlations, which arise after fixing *γ* due to the finite number of SNPs, would be included. Thus, 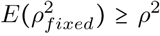 (with equality when *ρ*^2^ = 1). In contrast, SNP-heritability can be defined as 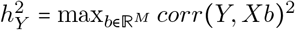 because the argmax equals the true causal effect size vector.

Mediation is not the same as causality; for instance, expression of a gene in one tissue might act as a proxy for expression of that gene in different, truly causal cell type and context. However, cis-genetic correlations are unlikely to reflect *reverse* causality (effects of trait on expression), as we can obtain an upper bound on the contribution of reverse causality to *ρ*^2^ for a polygenic trait (see Supplementary Note).

### Estimation using GECS regression

In order to estimate *ρ^2^* , we exploit a relation between the covariance between SNP effect sizes on expression and the covariance between SNP effect sizes on trait. This approach is related to Haseman-Elston Regression [32,33] (see Supplementary Note). If a SNP has a causal effect *β_i_* on the expression level of a gene, and that gene has an effect *α* on the trait, then the effect of the SNP on the trait is:

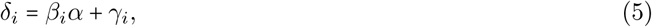

where *γ_i_* is the direct (non-mediated) effect. Now, given the eQTL effects of two SNPs, we have:

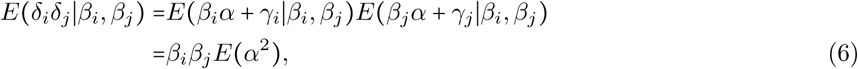

where we have assumed that *α* is independent of *β_i_*, *β_j_*; this assumption is discussed below. Under the independence assumption, *E*(*α*^2^) is proportional to *ρ*^2^ (see Supplementary Note). Replacing causal effects with marginal effects and allowing for multiple genes, we obtain (for SNPs not in LD):

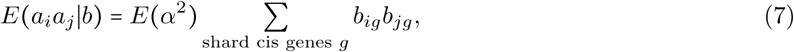

where *a_i_* is the marginal effect of SNP *i* on *Y* (including both mediated and non-mediated effects), and *b_ig_* is the marginal effect of SNP *i* on the expression level of gene *g.* By relating the average gene effect size, *E*(*α^2^*), with the total amount of mediation, *ρ*^2^, we recover equation (1); for a full derivation, see the Supplementary Note.

Motivated by equation (1), we estimate *ρ^2^* by regressing products of trait effect sizes on sums over genes of the products of eQTL effect sizes. The GECS regression estimate is equal to the regression coefficient times the mean expression heritability, divided by the trait heritability.

#### Key Assumptions

GECS regression requires certain assumptions that, if they were violated, would lead to a biased estimate of *ρ^2^*.

- **No LD or shared LD between SNP pairs** GE co-score regression is biased in the presence of LD or shared LD between SNP pairs. LD between SNP pairs induces a correlation between the noise terms in the effect size estimates for these SNPs (both the effects on expression and the effects on disease). This phenomenon leads to inflation in both the GE coscore regression numerator and the GE coscore regression denominator, and the magnitude of this effect is sample size dependent. Therefore, when the eQTL cohort is much smaller than the GWAS cohort, inadequate LD pruning can result in *downward* rather than upward bias in the estimate of *ρ*^2^, if the GE coscore regression denominator is more inflated than the numerator. This accounts for the increase in estimates of *ρ*^2^ when comparing 100% of SNPs retained in Figure 1-b with 95% of SNPs retained at a true *ρ*^2^ value of 0.5. Shared LD refers to LD with a third SNP; one measure of shared LD is *Σ_k_*(*r_ik_r_jk_*)^2^. In principle, SNPs with no LD could nonetheless have shared LD. In the presence of shared LD between *i, j* and *k*, the marginal effects of SNPs *i* and *j* contain a contribution from the causal effect of SNP *k*, in proportion to *r_ik_* and *r_jk_* respectively. This phenomenon occurs both for effects on trait and for effects on expression, regardless of whether the effects on trait are mediated by the effects on expression. Therefore, 
equation (1) requires that we exclude SNP pairs with shared LD. In practice, it is sufficient that SNPs have only a small amount of shared LD.
- **Independence between cis-genetic architecture and effects of expression on trait** In the derivation of equation (1) (see Supplementary Note), we assume that effects of gene expression on trait are drawn independently of eQTL effects: that is, genes with a certain type of genetic architecture do not have larger or smaller phenotypic effects on average. This assumption is needed because GECS regression implicitly weights some genes in the regression more heavily than others (that is, some genes contribute more to the variance in GE co-scores). Ideally, each gene would be weighted in proportion to its heritability, and indeed, if every eQTL for a given gene is doubled, then that gene will be weighted twice as heavily. However, weighting also depends on the number of independent eQTLs for a gene. Genes with only one large eQTL, due to either one causal variant or several causal variants in LD with each other, receive little weight in the regression due to LD pruning. Genes with multiple independent eQTLs are weighted more heavily, and the regression implicitly extrapolates from these genes to single-eQTL genes; thus, the estimator is biased if implicitly up-weighted genes have larger or smaller effect sizes on average than down-weighted genes.

#### Additional details

- **GE coscores.** Estimated GE coscores *𝒢̂* were computed for each subsample and from the whole sample: *𝒢̂_ij_* = Σ*_g_b̂_ig_b̂_jg_*.
- **LD estimates from expression data.** In-sample LD was also measured for all regression SNPs sharing a cis gene, in each subsample, using PLINK [47]. For pruning, the shared LD between SNPs *i*,*j* was estimated as 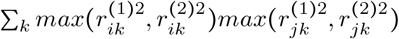, where 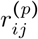 is the LD between SNP_S_ *i* and *j* in submale *p*. The LD between SNPS *i*, *j* was estimated as 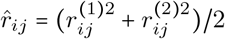.for discussion of the choice of estimator, see the Supplementary Note.
- **Conditioning on LD.** To remove effects from subtle LD below the threshold, we residualized estimate GE co-scores on estimated LD: 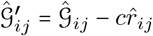.
- **Regression weights.** For each regression SNP, we computed the regression-SNP LD score, 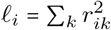, using in-sample LD values. For each SNP pair *i*, *j*, the regression weight was *w_ij_* = *1/*(*l_i,_ l_j_*).
- **LD pruning.** In order to ascertain SNP pairs having no shared LD and also no LD, SNP pairs having at least one shared cis gene were sorted by estimated shared LD and sorted by LD. The maximum of the two indices was then used to re-sort the SNP pairs, and the 25% of SNP pairs with highest combined LD were discarded (set *w_ij_ =* 0).
- **Regression numerator.** The regression numerator was:

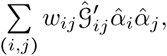
where *α̂_i_* is the estimated marginal effect size of SNP *i* on the trait and *𝒢̂_ij_* is the estimated GE coscore from the whole expression sample.
- **Regression denominator.** To eliminate attenuation bias, the regression denominator was not 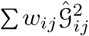 but rather
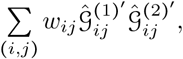
where 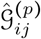 is the estimated GE coscore from subsample *p*.
- **Disease heritability estimation.** To estimate disease heritability and average cis heritability of gene expression levels, we use LD score regression [14,34].
- **Cis heritability estimation.** To estimate the average cis-heritability of gene expression levels, we apply cross-trait LD score regression [35] with no intercept to the subsampled summary statistics (treating expression in the two cohorts as separate traits); this procedure is similar to running cross-trait LD score regression with constrained intercept [35], except that it is robust to inflation or deflation in the LD score intercept arising from finite-sample effects. We use LD scores computed between cis regression SNPs and all reference SNPs within a 1cM window. In principle, it is preferable to compute LD scores between cis regression SNPs and cis reference SNPs only (see [37]), but in practice there is little difference (see Supplementary Note).
- **Block jackknife.** We estimated the standard error on the estimate of *ρ*^2^ by jackknifing on 50 blocks of contiguous SNPs, similar to LD score regression [34]. The same jackknife blocks were used for all traits. To estimate the standard error on the mean value of *ρ*^2^ across phenotypes, we jackknifed the mean. To assess significance of the estimates, we estimated the standard error of only the regression numerator (i.e., the covariance between GE coscores and products of trait *Z* scores), and applied a *Z* test to the regression numerator divided by its standard error.
- **Trait selection.** From the full set of 42 GWAS data sets (Table S2), pairs of data sets with estimated genetic correlation greater than 0.5 (estimated using cross-trait LD score regression [35]) were greedily pruned, beginning with the data sets having the highest genetic correlation. In each case, we retained the data set with higher power as measured by the significance of 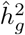 using LD score regression [34].

### Simulations

#### Primary simulation

For the primary simulation experiments, segments of chromosome 1 spanning about 1% of the genome were used, with cis windows of 601 SNPs (~ 2Mb), *M =* 5000 SNPs, and *G =* 60 partially heritable genes in *T =* 50 tissues (average 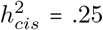, 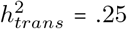). Sample size was N_eQTL_ = 1000 for the expression cohort, N_GWAS_ = 10,000 for the GWAS cohort, and N_reference_ = 500 for the reference panel cohort (used to compute LD scores); cohorts were disjoint. Entries of the M × GT eQTL effect size matrix were drawn initially from a point-normal distribution with on average 10 causal SNPs per cis locus and the same proportion of causal SNPs in trans. Effect sizes for cis SNPs were larger than effect sizes for trans SNPs, so that on expectation the sum of squared effect sizes within the cis region was equal to the sum of squared effect sizes in the trans region for each gene; as a result, on average half of gene expression heritability was in cis. To model correlations across tissues, the eQTL matrix was multiplied by a *GT* × *GT* matrix A, where *A_ii_ =* 1, *A_ij_ =* 0.8 if *i*, *j* correspond to the same gene in different tissues, and *A_ij_ =* 0 otherwise; afterwards, there was correlation 0.8 between expression levels of the same gene in different tissues. We drew a heritability value for each gene-tissue pair from a *Beta*(*3,* 3) distribution (which has mean 0.5). Then, the columns of the eQTL matrix were scaled so that the variance of the genetic component of each gene-tissue pair corresponded to these heritability values. Gaussian noise was added with covariance matrix *B*, where (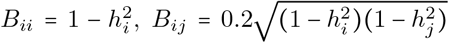) if i and j correspond to the same gene in different tissues, and *B_ij_* = 0 otherwise. Effect sizes of expression on trait were drawn from a point-Normal distribution (20% of genes causal), scaled so that the variance of the product of these effect sizes with the gene expression matrix corresponded to the simulated value of *ρ*^2^ (3). Effect sizes of SNPs directly on the trait were also drawn from a point-normal distribution, where the probability of SNP *i* being causal depended on its maximum eQTL effect size:

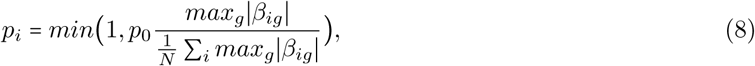

where we used *p*_0_ = 0.2. This results in a large amount of colocalization.

#### Secondary simulations

For simulations with different cis window sizes, we also modeled genetic correlations between adjacent genes (with overlapping but distinct cis windows) instead of across tissues. For simulations that included a comparison to the fixed-effect estimator, we used simulated, normally distributed genotypes with no LD. There were 500 SNPs and 100 genes, with disjoint cis windows of 5 SNPs per gene, and 100% of gene expression variance was determined by these 5 SNPs. Effects of expression on trait and of SNPs directly on trait were modeled similarly to the main simulations.

#### Fixed-effect estimator

Our fixed-effect estimator of *ρ*^2^ was the sum over genes of the squared estimated genetic covariance between the trait and the gene, divided by the total heritability; the genetic covariance was estimated as the sum over cis SNPs of the products of the estimated effect sizes on each trait. Because cis-heritability of gene expression was 1, we did not normalize by estimated expression heritability.

#### Partitioned LD score regression

We ran partitioned LD score regression [14] on the simulated data, using LD scores derived from 500 individuals (not overlapping with the GWAS or eQTL cohorts). SNPs were ranked by their maximum *χ^2^* statistic over cis genes, and the top 25% of SNPs were called as eQTLs. Then, we regressed trait *χ*^2^ statistics on the vector of LD scores as well as the binary vector of eQTL calls, with intercept fixed at 1. We estimated the standard error of these estimates using 20 jackknife blocks, and obtained p values using a *Z* test. This approach is similar, in practice, to the approach of computing LD scores to called eQTLs; however, since eQTLs are called based on marginal, rather than causal, effect sizes, the latter approach assigns overly large scores to regions with more LD, since SNPs in these regions are more likely to be called as eQTLs (however, both of these approaches produce similar results).

#### Standard error assessment

In order to assess the jackknife estimate of the standard errors, we computed point estimates of 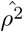 of *ρ^2^* as well as jackknife estimates *σ̂* of the standard deviation of the estimate. We computed normalized errors, 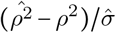, and reported the root mean squared normalized error. This metric enables us to determine whether error bars are expected to give accurate confidence intervals; if instead we computed the average estimated standard error with the empirical standard error, it is possible the estimated standard error would be unbiased, and yet confidence intervals derived from estimated standard errors would be badly miscalibrated.

### GTEx data set

For every gene and every tissue, effect size estimates *b̂* were produced for all regression SNPs within 1Mb of the TSS (G ~ 24,000 genes × *T =* 44 tissues; M ~ 170,000 regression SNPs), both for the full expression cohort and for two random subsamples containing half of the cohort. Pooling of tissues increases power (see Supplementary Note). We used FastQTL [45], restricted to unrelated individuals, and corrected for sex, genotyping platform, 3 PCs, and 15-35 PEER factors [46], consistent with GTEx consortium methods [3]. We additionally restricted to European samples.

## Supplementary Note

### Reverse causality

We can obtain an upper bound on the contribution of reverse causality to *ρ*^2^ for a polygenic trait. Related intuition is discussed in reference [24]. The worst-case hypothetical scenario is a fully-heritable trait T that explains the entire trans-genetic component of gene expression, in addition to some of the cis-genetic component for every gene. For gene g, the covariance between *Z̃_g_* and *T* is

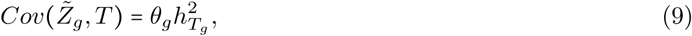
where *θ_g_* is the effect of T on *Z_g_* and 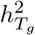 is the SNP-heritability of T localizing to the cis region of SNP *g*. Now, the marginal effect of gene *g* on *T* (due to reverse causality) is *Cov*(*Z̃_g_*,T)*/Var*(*Z̃*_g_), and

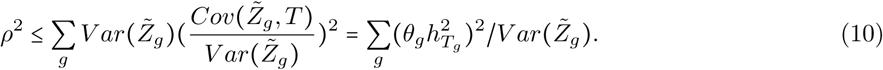

That is, if we regress *y* on a vector *X*, the variance explained is at most the sum of the squared marginal effect sizes, normalized by the variance in each *X_i_*. The reason this number cannot be large is that if *T* is a polygenic trait, then the trans-genetic component of *T* must explain much more than the cis-genetic component. If the heritability of *T* is spread evenly across the genome, then we have 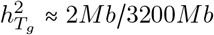. 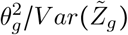 is at most the total heritability of gene *g* divided by its cis heritability; if we take 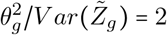 and *G =* 20,000 genes, then we obtain an upper bound on *ρ*^2^ of

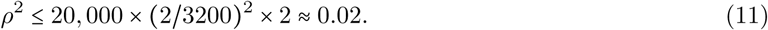

If we wish to bound the contribution of reverse causality to our overall results, rather than to an individual trait, then we can revise the assumption that T explains the entire trans-genetic component of gene expression; this study considered 30 genetically distinct traits, and it is impossible that all of them have this property. If instead each trait individually explains 1/30^th^ of this trans-genetic component, then we have 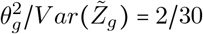, and we have *ρ*^2^ ≤ 0.001. This type of argument also extends to causal scenarios where a small number of intermediaries affect both the trait and gene expression. If there are k such intermediaries, then a similar bound is obtained for *ρ*^2^, differing by a factor of *k*.

### Alternate approaches

We considered several alternative approaches, but these would not produce reliable estimates of *ρ*^2^, because they fail to distinguish between mediation and colocalization.

- **Comparison of** χ^2^ **statistics.** One possible approach is to regress phenotypic χ^2^ statistics on sums of eQTL χ^2^ statistics, possibly with an LD score covariate. This would be an attractive method for measuring colocalization, and under the assumption that all colocalization is driven by mediation, it would provide an estimate of *ρ*^2^.
- **Aggregation of fixed-effect estimates.** Suppose we were given a vector s of fixed effect cis-genetic correlations, i.e. for gene *k*, *s_k_* = *Corr*(*Xβ_k_*,*Y*). A natural approach would be to aggregate these estimates; for example, in the absence of cis-genetic correlations between genes, an estimator of *ρ*^2^ would be 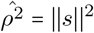. This estimator is used in Figure 1-c. However, this estimator ignores the difference between the fixed-effect quantity *s* and the random-effect quantity *α*. Moreover, the difference between ||*s*||2 and ||*α*||^2^ depends on the amount of colocalization between eQTLs and non-mediated effects, with a larger difference in the presence of stronger colocalization; thus, this approach also fails to distinguish between mediation and colocalization.
- **Imputation of a gene expression relatedness matrix.** A third approach is to impute a gene expression relatedness matrix (“GERM”) using existing methods [23, 24], which could be used as the input to a standard heritability estimator [31,33,48]. Unfortunately, the GERM will “tag” the genetic relatedness matrix (GRM), leading to upward bias in the estimate. Even if the GRM is used as a covariate, colocalization will allow the GERM to tag *important* genetic variants, and there will still be inflation.

### Derivation of moment condition

Here, we derive equation (1) under the assumption that gene effect sizes are drawn independently of the eQTL entries *β* and the LD matrix *R*:

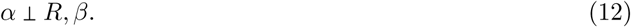

Violations of this assumption are discussed below. The marginal effect of a SNP on the phenotype is

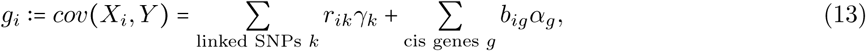

where *B = Rβ =* (*b_ig_*) is the marginal eQTL matrix. The covariance between marginal SNP effect sizes *g* is

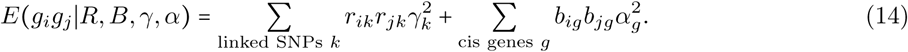

Now we marginalize out *γ* and *α*, using the fact that *γ* is uncorrelated (on expectation) with *β*:

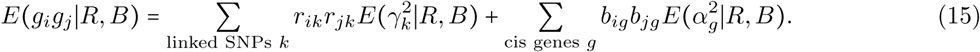

Replacing true effect sizes with their estimates from a finite sample, we have

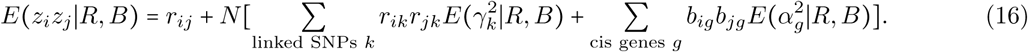

We wish to avoid assuming that (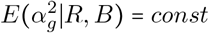), i.e. that non-mediated effects do not colocalize with eQTLs. Therefore, we remove pairs (*i, j*) of linked SNPs, so that *r_ik_ r_jk_* ≈ 0. The term *r_ij_* also vanishes. By assumption (12), 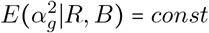. Defining 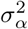 as this constant, we have that

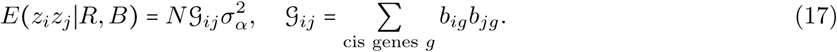

The independence assumption (12) also implies that

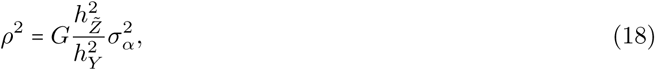

and we recover (1).

Some types of violations of the independence assumption, but not others, lead to violations of (1) and to bias in the estimator. Only genes with more than one independent eQTL contribute to *𝒢*, so if genes with more polygenic cis-genetic architectures have larger or smaller effect sizes on average, the moment condition is violated and bias is introduced in the estimator. In effect, GECS regression extrapolates from genes with multiple eQTLs to genes with only one eQTL. In contrast, violations of (12) mediated by differences in cis heritability do not lead to violations of (1), even though they do lead to violations of both (16) and (18); this is because more-heritable genes contribute more heavily to *𝒢*, and they contribute proportionally to *ρ^2^.* This can be seen by considering the effect of adding a gene with zero cis heritability. Similarly, if cis-genetically correlated genes have correlated effect sizes, it leads to compensatory violations of (16) and (18), but not (1). This can be seen by considering the effect on of adding a perfectly-correlated copy of every gene in the genome, with an identical effect size to its original, while halving all effect sizes.

### Power and number of tissues

GECS regression improves power by concatenating tissues, treating the same gene in *k* tissues as *k* distinct genes. When cis-genetic correlations across tissues are strong and environmental correlations are weak, this approach increases power by increasing the effective heritability of each gene. In particular, if the tissues are perfectly cis-genetically correlated, and there is no other covariance between the tissues (due to environment or trans-genetic correlations), then the effective cis-heritability is

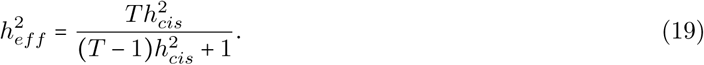

To see this, it is useful to construct a vector *b^*^ = R̂β,* where *R̂* is the in-sample LD matrix and *β* is the *true* causal effect size vector. Note that b^*^ differs from both the true marginal effect size vector *b* = *Rβ* and the estimated marginal effect size vector *b̂*. *b*^*^ has the property that in the absence of environmental noise 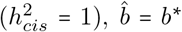. Similarly, given a large number of perfectly cis-genetically correlated tissues from each individual in the eQTL cohort, the average across tissues of the estimated effect size vectors converges to b^*^. We can write the estimated effect vector for SNP *i* and tissue *t* as 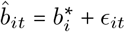, where

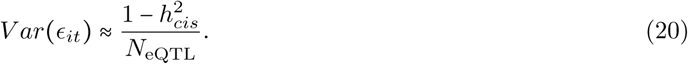

As a proxy for GECS regression power, we can use

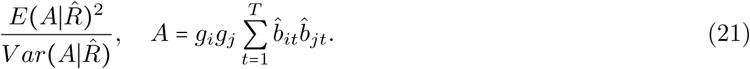

where *g_i_* is the effect size of SNP *i* on the phenotype; we use *g_i_* instead of *ĝ_i_* to isolate the component of power that is dependent on the eQTL cohort (i.e., when the GWAS cohort is large). We further condition on the in-sample eQTL cohort LD matrix *R̂* in order to isolate the component of power that is determined by the number of tissues and by the cis-heritability.

First,

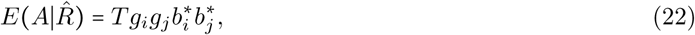

where we have used that *i* and *j* are not in LD (so *ϵ_it_* is uncorrelated with *ϵ_jt_*). Next,

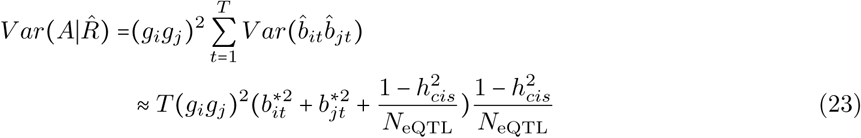

where we have used independence of noise terms across tissues in the first line and equation (20) together with the assumption of no LD between SNPs *i* and *j* in the second line. Disregarding terms of size 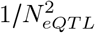 and assuming *g_i_ g_j_ ≠* 0 and 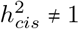, we have

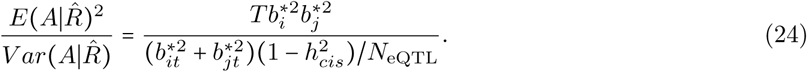

Taking 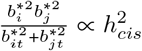, we have

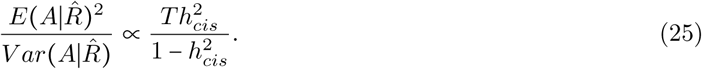

Finally, we solve

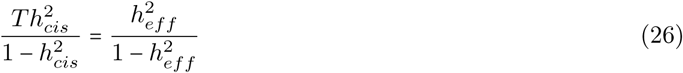

to obtain equation (19).

### Additional sources of bias

#### Bias in GECS regression due to expression heritability flanking the cis window

GECS regression only considers cis-eQTL effects, defined using some cis radius. If the chosen cis radius is too small, the primary effect is that mediation will be missed for variants outside of the window (Figure S2). However, there is also a competing effect, which occurs as a result of LD between cis SNPs in conjunction with genetic correlations between genes. To illustrate this effect, consider an extreme example: the genome comprises 2 perfectly correlated genes and 20 SNPs, with SNPs 1, 2,…, 10 being in cis for gene A and SNPs 11, 12, …20 being in cis for gene B. SNP 1 is in perfect LD with SNP 11, SNP 2 is in perfect LD with SNP 12, and so on. If we apply GECS regression, the estimator will be upwardly biased by a factor of 2, since genetic effects on gene A perfectly tag effects on gene B, and this will be reflected in the GECS regression numerator but not in the GECS regression denominator (the variance in GE coscores). In contrast, if the cis radius is extended so that every SNP is cis for both genes, then there is no longer any bias, since the tagging is also reflected in the GECS regression denominator. Thus, this type of bias requires genetic correlations between genes with cis windows that do not perfectly overlap (i.e., not only between the same gene in different tissues), and it also requires LD between SNPs that are in cis for one gene or the other but not both. It is expected to disappear when the cis window is sufficiently large, as SNPs on the boundaries of the cis region (which might have LD with SNPs outside the cis region) will explain little expression heritability.

#### Bias in cis-heritability estimation due to LD between cis and trans SNPs

We estimate cis-heritability of gene expression levels using LD score regression on cis SNPs with LD scores computed to both cis and trans SNPs. In principle, it is preferable to use LD scores between regression SNPs and cis reference SNPs and trans reference SNPs separately [37]; the inclusion of non-cis reference SNPs introduces a small amount of downward bias in the cis-heritability estimates. If SNPs flanking the cis region explain zero heritability, then estimates are biased toward zero by a factor of approximately *ℓ̅_cis_/ℓ̅,* where *ℓ̅_cis_* is the average LD score between regression SNPs and cis reference SNPs, and *ℓ̅* is the average LD score between regression SNPs and all reference SNPs. For a cis window size of ±1Mb, this bias is small (at most a few percent), since cis SNPs have more LD with other cis SNPs than with trans SNPs; it is more substantial when a smaller cis window size is used.

### Relationship with Haseman-Elston Regression

GECS regression regression is related to Haseman-Elston (HE) Regression [32,33]. HE regression relates the covariance between the phenotypes of two individuals with the covariance between their genotype vectors (i.e., their genetic relatedness):

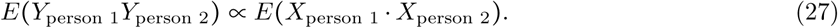

This follows from

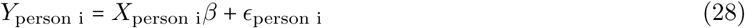

where *β* is random and, crucially, E(*ϵ*_1_*ϵ*_2_) = 0. Similarly, GECS regression relates the covariance between the phenotypic *effects* of two SNPs with the covariance between their eQTL vectors (i.e., the GE coscore):

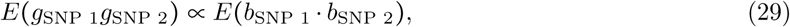

and this follows from

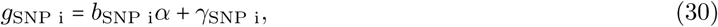

where E(γ_1_ γ_2_) = 0 specifically for SNPs with no shared LD.

### Choice of LD estimator

We estimate LD using in-sample LD for the two subsamples that are used to estimate the GE coscore regression denominator. We have found that when out-of-sample LD is used, estimates are inflated due to residual LD. It is sufficient to remove in-sample LD for the eQTL cohort only (and not the GWAS cohort) because even if there is residual LD in the GWAS cohort (owing to LD mismatch or finite sample size), this residual LD will not correlate with residual LD in the eQTL cohort. It is necessary to use in-sample LD for the cross-validation subsamples, since if we ascertain SNP pairs with low total LD, then this ascertainment procedure introduces dependencies between the cross-validation subsamples, leading to downward bias in the GE coscore regression denominator.

### Supplementary Tables

Table S1: Performance of GE score regression in simulations using real genotypes and simulated gene expression and phenotypes (see Excel file). Simulations included strong colocalization between eQTLs and non-mediated disease SNPs. (a) Estimates were approximately unbiased when 75% of SNP pairs were retained. (b) Estimates were inflated when no LD pruning was applied, but LD pruning reduced the bias. Dashed horizontal lines indicate true values of *ρ*^2^. (c) Estimates of *ρ*^2^ using GECS regression and a naive fixed-effect estimator at different eQTL and GWAS sample sizes, in a simplified simulation setup including colocalization but no mediation. (d) Comparison of power to detect an effect between GECS regression and partitioned LD score regression, using eQTLs as an annotation.

**Table S2:** GWAS datasets analyzed. The first 30 phenotypes (above horizontal line) were included in the primary analysis, and the remaining phenotypes were excluded from the primary analysis due to high genetic correlation. Of the first 30 phenotypes, related traits are grouped and order is alphabetical within groups. To avoid bias due to regression SNP selection, GWAS using specialty chip data were excluded, similar to [14].

| Phenotype | Reference / Dataset | N |
| --- | --- | --- |
| Anorexia | Boraska et al., 2014 Mol Psych | 32143 |
| Autism Spectrum | PGC Cross-Disorder Group, 2013 Lancet | 10263 |
| Putamen volume | Franke et al., 2016 Nat Genet | 11840 |
| Neuroticism | Okbay et al., 2016 Nat Genet | 170911 |
| Schizophrenia | SCZ Working Group of the PGC, 2014 Nature | 70100 |
| Years of Education | Okbay et al., 2016 Nature | 328917 |
| Waist-hip ratio (BMI adjusted) | UK Biobank | 145375 |
| HDL | Teslovich et al., 2010 Nature | 97750 |
| LDL | Teslovich et al., 2010 Nature | 93354 |
| Triglycerides | Teslovich et al., 2010 Nature | 94461 |
| Smoking status | UK Biobank | 145227 |
| Blood pressure (Systolic) | UK Biobank | 134011 |
| HbA1C | Soranzo et al., 2010 Diabetes | 46368 |
| Type 2 Diabetes | Morris et al., 2012 Nat Genet | 60786 |
| Celiac Disease | Dubois et al., 2010 Nat Genet | 15283 |
| Crohn's Disease | Jostins et al., 2012 Nature | 20883 |
| Lupus | Bentham et al., 2015 Nat Genet | 14267 |
| Multiple sclerosis | IMS Genetics Consortium, 2011 Nature | 27148 |
| Primary Biliary Cirrhosis | Cordell et al., 2015 Nat Commun | 13239 |
| Rheumatoid Arthritis | Okada et al., 2014 Nature | 37681 |
| Ulcerative Colitis | Jostins et al., 2012 Nature | 27432 |
| Asthma | UK Biobank | 145416 |
| Eczema | UK Biobank | 145416 |
| Age at Menarche | UK Biobank | 74944 |
| Age at Menopause | UK Biobank | 44410 |
| Lung FEV1/FVC ratio | UK Biobank | 123935 |
| Lung Forced Expiratory Volume (FVC) | UK Biobank | 123935 |
| Bone mineral density (heel) | UK Biobank | 141441 |
| BMI | Speliotes et al., 2010 Nat Genet | 122033 |
| Height | Lango Allen et al., 2010 Nature | 131547 |
| Bipolar Disorder | BIP Working Group of the PGC, 2011 Nat Genet | 16731 |
| Blood pressure (Diastolic) | UK Biobank | 134011 |
| BMI | UK Biobank | 145209 |
| College Education | UK Biobank | 144204 |
| Coronary Artery Disease | Schunkert et al., 2011 Nat Genet | 77210 |
| Depressive symptoms | Okbay et al., 2016 Nat Genet | 161460 |
| Ever Smoked | TAG Consortium, 2010 Nat Genet | 74035 |
| Fasting Glucose | Manning et. al., 2012 Nat Genet | 58074 |
| Height | UK Biobank | 145368 |
| Hypertension | UK Biobank | 145379 |
| Subjective wellbeing | Okbay et al., 2016 Nat Genet | 298420 |
| Years of Education | Rietveld et al., 2013 Science | 126559 |

**Table S3:** GTEx tissues included in the analysis, and sample size. Tissues marked with an asterisk were included in the brain and brain-related tissues analysis (Table 3).

| Tissue | $N_{\text{eQTL}}$ |
| --- | --- |
| Adipose Subcutaneous | 298 |
| Adipose Visceral | 185 |
| Adrenal Gland | 126 |
| Artery Aorta | 197 |
| Artery Coronary | 118 |
| Artery Tibial | 285 |
| Brain Anterior* | 72 |
| Brain Caudate* | 100 |
| Brain Cerebellar* | 89 |
| Brain Cerebellum* | 103 |
| Brain Cortex* | 96 |
| Brain Frontal* | 92 |
| Brain Hippocampus* | 81 |
| Brain Hypothalamus* | 81 |
| Brain Nucleus* | 93 |
| Brain Putamen* | 82 |
| Breast Mammary | 183 |
| Cells EBV-transformed | 114 |
| Cells Transformed | 272 |
| Colon Sigmoid | 124 |
| Colon Transverse | 169 |
| Esophagus Gastroesophageal | 127 |
| Esophagus Mucosa | 241 |
| Esophagus Muscularis | 218 |
| Heart Atrial | 159 |
| Heart Left | 190 |
| Liver | 97 |
| Lung | 278 |
| Muscle Skeletal | 361 |
| Nerve Tibial* | 256 |
| Ovary | 85 |
| Pancreas | 149 |
| Pituitary* | 87 |
| Prostate | 87 |
| Skin Not | 196 |
| Skin Sun | 302 |
| Small Intestine | 77 |
| Spleen | 89 |
| Stomach | 170 |
| Testis | 157 |
| Thyroid* | 278 |
| Uterus | 70 |
| Vagina | 79 |
| Whole Blood | 338 |

**Table S4:** Estimated heritability mediated by expression for pairs of genetically correlated traits or studies. P values were obtained by jackknifing on the difference between the estimates.

| Phenotype 1 | $\rho^2$ | std err | Phenotype 2 | $\rho^2$ | std err | $p_{\text{difference}}$ | genet corr | std err |
| --- | --- | --- | --- | --- | --- | --- | --- | --- |
| Depressive symptoms | 0.17 | 0.07 | Neuroticism | 0.12 | 0.02 | 0.03 | 0.78 | 0.05 |
| Depressive symptoms | 0.17 | 0.07 | Subjective wellbeing | 0.35 | 0.34 | 0.16 | -1.18 | 0.14 |
| Neuroticism | 0.12 | 0.03 | Subjective wellbeing | 0.35 | 0.34 | 0.07 | -0.99 | 0.09 |
| Bipolar Disorder | 0.25 | 0.09 | Schizophrenia | 0.12 | 0.02 | 0.02 | 0.79 | 0.06 |
| Coronary Artery Disease | 0.77 | 0.66 | Hypertension | 0.15 | 0.05 | 0.33 | 0.94 | 0.24 |
| Diastolic | 0.13 | 0.03 | Hypertension | 0.15 | 0.05 | 0.03 | 0.74 | 0.03 |
| Ever smoked | 0.21 | 0.1 | Smoking status | 0.13 | 0.02 | 0.1 | 0.96 | 0.09 |
| Diastolic | 0.13 | 0.03 | Systolic | 0.12 | 0.04 | 0.81 | 0.75 | 0.03 |
| Hypertension | 0.15 | 0.05 | Systolic | 0.12 | 0.04 | 0.05 | 0.72 | 0.03 |
| Fasting Glucose | 0.24 | 0.2 | Type 2 Diabetes | 0.27 | 0.1 | 0.43 | 0.74 | 0.14 |
| Years of Ed. - Okbay | 0.09 | 0.02 | Years of Ed. - Rietveld | 0.2 | 0.04 | 0.002 | 1.07 | 0.03 |
| BMI - Speliotes | 0.1 | 0.02 | BMI - UKB | 0.1 | 0.03 | 0.59 | 0.97 | 0.03 |
| Years of Ed. - Okbay | 0.09 | 0.02 | College Ed. - UKB | 0.12 | 0.02 | 0.04 | 0.97 | 0.02 |
| Years of Ed. - Rietveld | 0.2 | 0.04 | College Ed. - UKB | 0.12 | 0.02 | 0.14 | 0.95 | 0.05 |
| Height - Allen | 0.13 | 0.03 | Height - UKB | 0.12 | 0.03 | 0.92 | 1.02 | 0.02 |

Table S5: Calibration of jackknife standard errors (see Excel file). Signed empirical errors were divided by the jackknife standard errors, and the standard deviation of the normalized errors is reported for different values of *ρ*^2^.

Table S6: Effect of cis window size on estimated *ρ*^2^ in simulations (see Excel file). (a) Estimates of *ρ*^2^ for different window sizes, either in the presence of genetic correlations across tissues or in the presence of genetic correlations between adjacent genes. (b) Estimates of *ρ*^2^ per unit of cis heritability, either in the presence of genetic correlations across tissues or in the presence of genetic correlations between adjacent genes.

Table S7: Stability of estimated *ρ*^2^ across LD thresholds (see Excel file). Estimates of *ρ*^2^ were obtained for different LD thresholds and averaged across phenotypes. Increasingly stringent thresholds were placed on the amount of shared LD for SNP pairs in the regression, and standard errors were calculated by jackknifing on the mean.

Table S8: Effect of cis window size on estimated *ρ*^2^ (see Excel file). (a) Heritability mediated by expression was variable for small window size values, but stable for larger window sizes. (b) Heritability mediated by expression, per unit of expression heritability, was larger when the cis window size was smaller than ±1Mb.

Table S9: Dependence on number of tissues (see Excel file). Estimates of *ρ*^2^ were obtained using random samples of tissues and averaged across non-redundant phenotypes. Error bars indicate jackknife standard errors (which do not include variability due to true differences among tissues).

### Supplementary Figures

**Figure S1:**
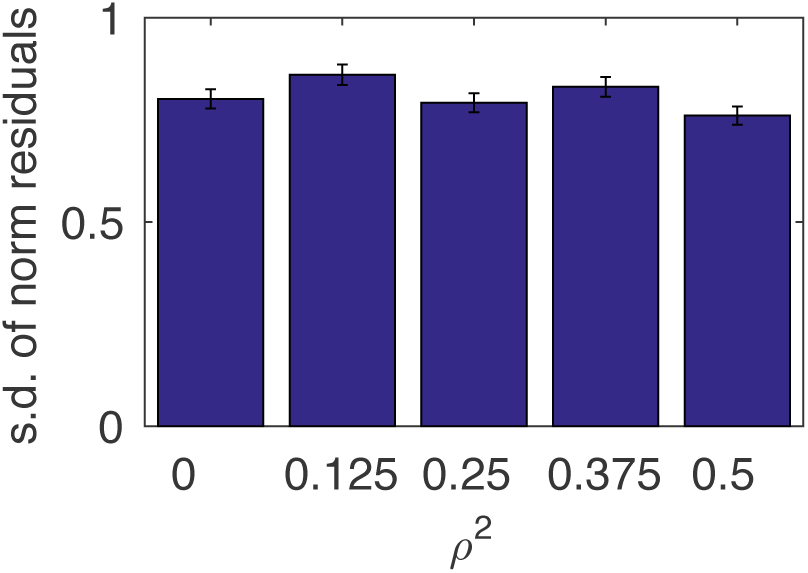
Calibration of jackknife standard errors. Signed empirical errors were divided by the jackknife standard errors, and the standard deviation of the normalized errors is reported for different values of *ρ*^2^. Error bars indicate standard errors on the sample standard deviation, based on 288 simulations. Numerical results are reported in Table.

**Figure S2:**
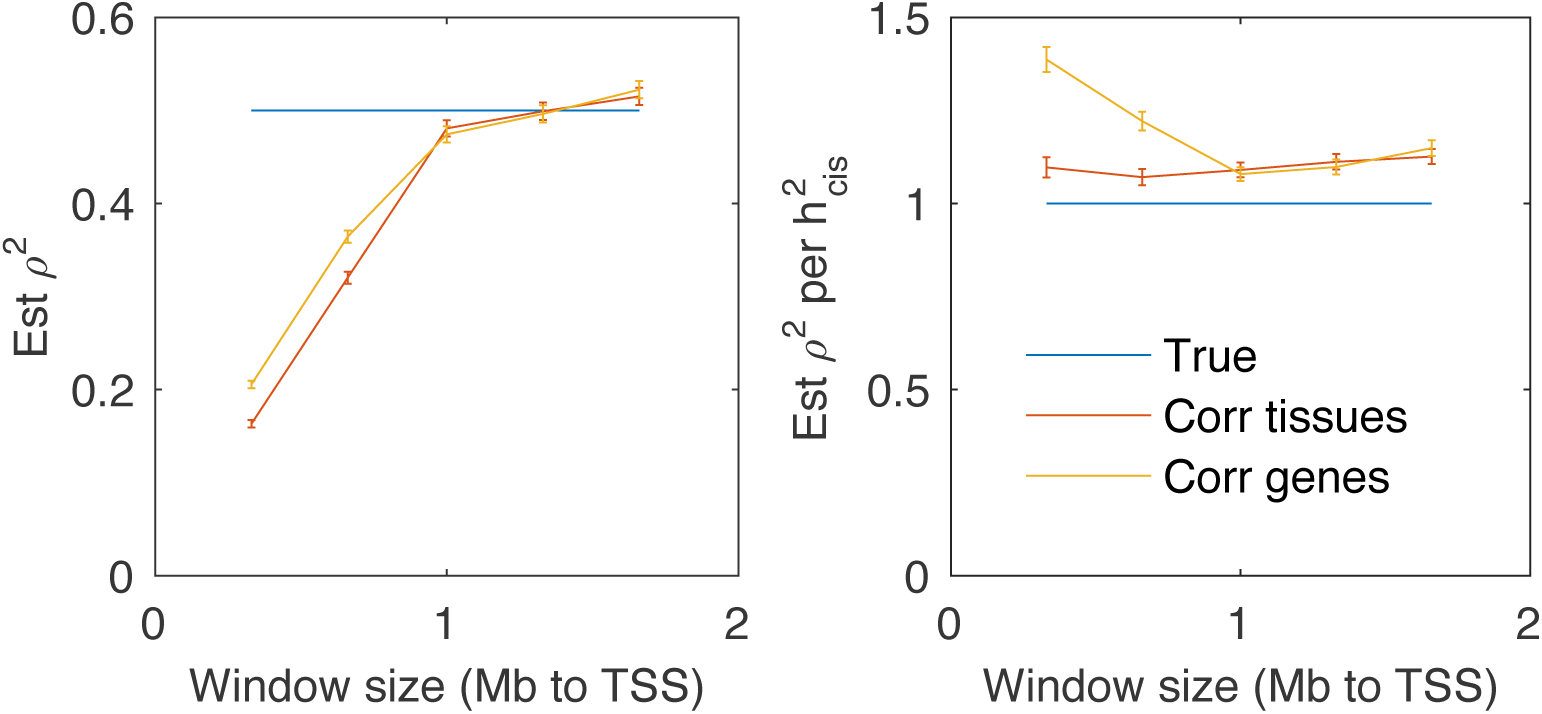
Effect of cis window size on estimated *ρ*^2^ in simulations. (a) Estimates of *ρ*^2^ for different window sizes, either in the presence of genetic correlations across tissues or in the presence of genetic correlations between adjacent genes. (b) Estimates of *ρ*^2^ per unit of cis heritability, either in the presence of genetic correlations across tissues or in the presence of genetic correlations between adjacent genes. Error bars indicate standard error based on 576 simulations. Numerical results are reported in Table S6.

**Figure S3:**
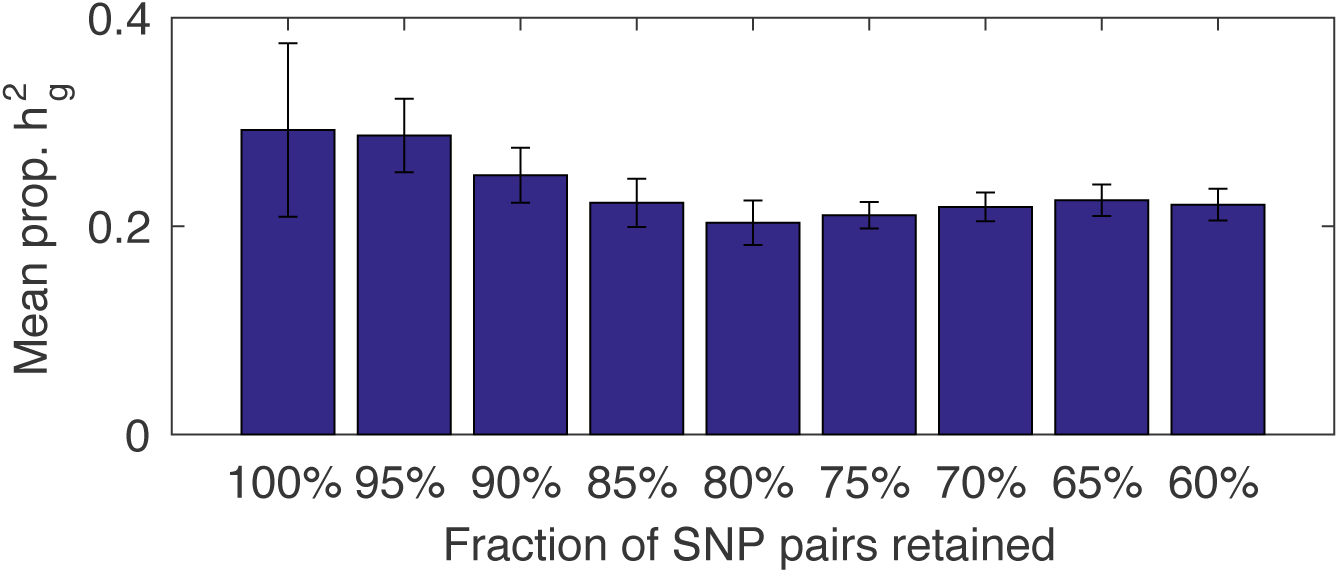
Stability of estimated *ρ*^2^ across LD thresholds. Estimates of *ρ*^2^ were obtained for different LD thresholds and averaged across phenotypes. Increasingly stringent thresholds were placed on the amount of shared LD for SNP pairs in the regression, and standard errors were calculated by jackknifing on the mean. Numerical results are reported in Table S7.

**Figure S4:**
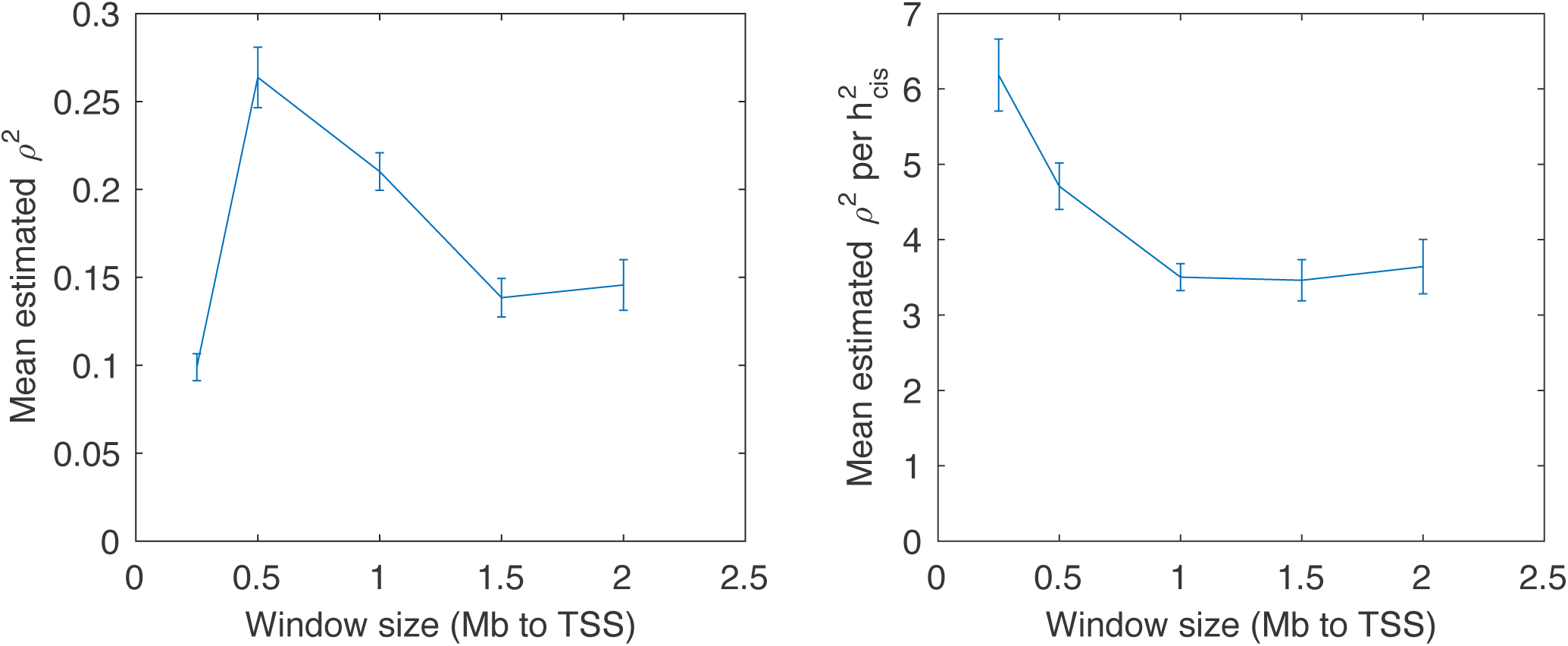
*ρ^2^.* (a) Heritability mediated by expression was variable for small window size values, but stable for larger window sizes. (b) Heritability mediated by expression, per unit of expression heritability, was larger when the cis window size was smaller than ±1Mb. Numerical results are reported in Table S8.

**Figure S5:**
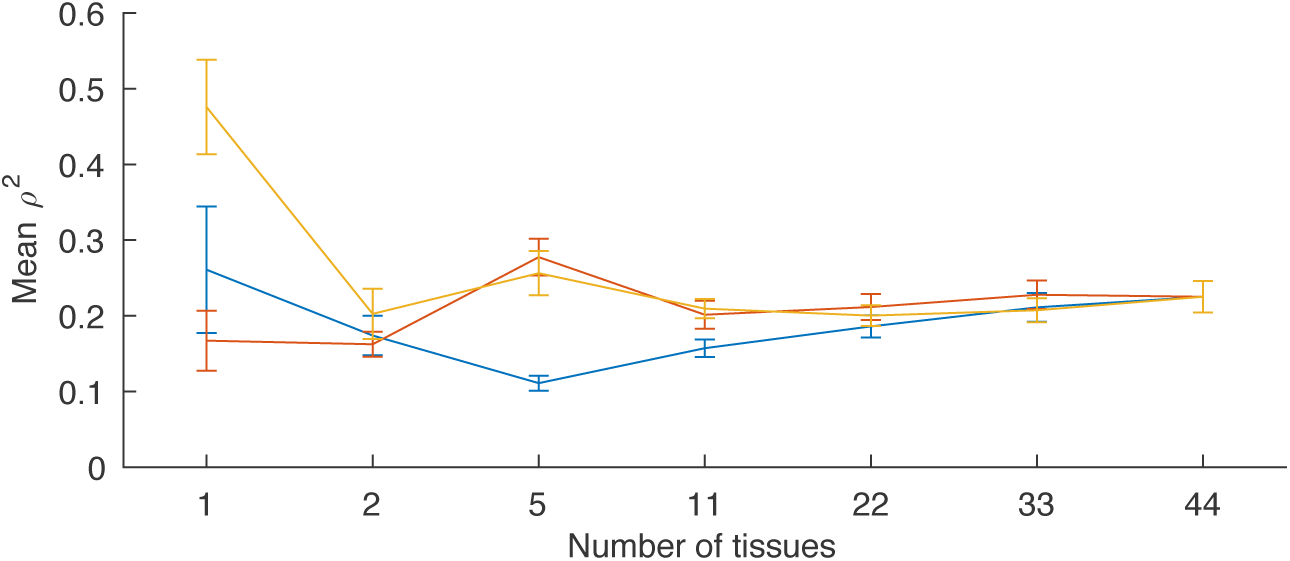
Dependence on number of tissues. Estimates of *ρ^2^* were obtained using random samples of tissues and averaged across non-redundant phenotypes. Error bars indicate jackknife standard errors (which do not include variability due to true differences among tissues). Numerical results are reported in Table S9.

